# Socioeconomic factors associated with dysentery in children under-five years from developing countries

**DOI:** 10.1101/664607

**Authors:** Ángela María Pinzón-Rondón, Carol Jisseth Zarate-Ardila, Laura Parra-Correa, Alisson Zarate-Ardila, Paola Lozada-Calderón, Leire Di Cecco

**Affiliations:** Escuela de Medicina y Ciencias de la Salud, Universidad del Rosario, Bogotá, Colombia; Facultad de Medicina y Ciencias de la Salud, Universidad Militar Nueva Granada, Bogotá, Colombia

**Keywords:** Dysentery, Multilevel Analysis, risk factors, prevalence

## Abstract

**Objective:** Dysentery represents 10% of all causes of acute diarrhea in the world and recognizing the implied proximal and distal social factors at different levels would impact on every related outcome. Our purpose is to identify mother, household and country characteristics that favor the presence of dysentery in children under 5 years old.

**Methods:** We conducted a multilevel analysis of data from phase V of the Demographic and Health Survey and the World Bank, which included 38,762 children from 33 countries.

**Results:** Prevalence of dysentery was 14.74%. GDP per-capita was negative associated (OR= 0.75; 95% CI 0.71-0.78) and Gini index was positive associated (OR= 1.23; 95% CI 1.19-1.28). Additionally, child age (OR= 0.99; 95% CI 0.99-1.00), mother age (OR= 1.01; 95% CI 1.00-1.01), employed mother (OR= 1.11; 95% CI 1.02-1.20), and number of household members (OR= 1.02; 95% CI 1.01-1.03) have significant positive associations with the presence of dysentery, while complete immunization schedule (OR= 0.88; 95% CI 0.81-0.96), duration of breastfeeding (OR= 0.81; 95% CI 0.75-0.89), and type of residence (OR= 0.87; 95% CI 0.79-0.97) have significant negative associations with having the illness. Finally, each of the categories of wealth index showed a significant association with dysentery (p-value < 0.001).

**Conclusions:** Lower per capita GDP and higher Gini coefficient are associated with the development of dysentery, regardless of characteristics of children, their mother, and household. Future and present public health programs should address these issues in order to impact on the occurrence of this illness.

**Author summary:** Dysentery represents 10% of all causes of acute diarrheal disease. Diarrhea is the fifth cause of worldwide death in children under five years old. It is particularly important to assess and prevent this condition because the early years of life are critical since it is the period when the brain develops most rapidly and has a high capacity for change. Complications associated with dysentery such as malnutrition and convulsive episodes could have a negative effect in this aspect.

Our purpose is to identify the country proximal and distal socioeconomic factors that favor the presence of dysentery in children under five years old from low and middle-income developing countries in order to impact on the occurrence of this illness and its related outcomes. Studying associated factors with developing dysentery during an episode of acute diarrhea could be the base upon which we can diminish mortality from this illness through national policies to impact on national, community and household aspects.

## Introduction

Acute diarrheal disease (ADD) is defined as the presence of three or more abnormally loose or watery stools in 24 hours. It is the second cause of death in children under five years old [1,2], with a worldwide prevalence of about 8% [3]. This illness is caused usually by the presence of microorganisms in contaminated food or drinking water [4], most of the times by rotavirus, which accounts 527,000 deaths annually. Eighty-two percent of these deaths occur in the poorest countries [5]. The second leading cause of ADD is related to a bacterial infection, in which *E. coli enteropathogenic* is the most important involved microorganism [6].

Dysentery is an intestinal inflammation that can lead to severe diarrhea with mucus or blood in the stools. It is a particularly worrying presentation of ADD [1] usually accompanied by fever, abdominal pain and impaired general conditions. Hence, it is defined as the presence of macroscopic blood and mucus in the stools [1,4]

According to the World Health Organization (WHO), dysentery represents 10% of all causes of ADD and accounts for 15% of all deaths from this cause. It is also the fifth cause of death worldwide in children under five years old [7]. In developing countries, shigellosis is the most frequent cause of dysentery in children, with 99% of all cases in comparison to developed countries whose main etiology is seasonal virus [4,6,8]. Complications associated with dysentery goes from dehydration, malnutrition, and cognitive impairment to severe outcomes as convulsive episodes, uremic syndrome and death [8]. It is worth recognizing, that children are at higher risk to acquire this mentioned infection, not only because they have more contact with soil, but also because their immune system is still immature for establishing an appropriate response to any microorganism [2].

In a recent study, we evaluated the relationship between some characteristics of developing countries and the occurrence of diarrheal disease. We concluded that residents of nations with higher inequality and lower incomes have greater probabilities of having diarrhea, especially when there is a lack of household wealth and mother’s education [9].

Other factors that showed strong positive association with diarrhea were female sex of the child, younger age of the child, incomplete immunization status, birth weight, lack of education of the mother and an unemployed mother [9].

Recognizing dysentery risk factors would reduce not only mortality rates in children under five years old but also would impact in every outcome related to this disease. The aim of this article is to identify mother, household and country characteristics that favor the presence or absence of dysentery in children under five years old by analyzing the Demographic and Health Survey (DHS) phase V in 33 nations.

## Methods

This is a cross-sectional, transnational and multilevel study. We used level-1 data (child, mother and household characteristics) from the Demographic and Health Survey (DHS) phase-V [10] and level-2 data (country characteristics) from the World Bank (WB) country data [11].

The DHS phase-V collected data from 41 developing countries from 2004 to 2010. A nationally representative, probabilistic sample including rural and urban areas was collected from each participating country. Respondents were selected through a multistage, stratified sampling procedure of households. Between 5,000 and 30,000 households were surveyed per country. Data was gathered for the countries included in Attachment 1. We excluded Ukraine from the analysis because this country did not apply the child health module of the survey. Information from the remaining 40 countries was merged to create a single dataset, which included 395,485 households with children. The dataset was further limited to biological mothers answering the survey to assure comparability (384,662), living children (359,527), permanent household residents (349,849) and cases with complete information in variable diarrhea (348,706). Afterward, the database was limited to cases that reported diarrhea during the two weeks before the interview (49,065) in order to know how many of them had dysentery. Then, the database was finally limited to cases with complete information in the variable dysentery (38,762). Finally, we have information from 33 countries because Bangladesh, Benin, Congo, Democratic Republic of the Congo, Indonesia, Mali, and Niger did not have information about dysentery [10].

After careful analysis, we concluded that the WB country data was the best source of level-2 data in this study because of its country comparability and robustness when compared to data from other sources. These data included the 2010 indicators: per capita gross domestic product (GDP), Gini-coefficient, and health expenditure as a percentage of GDP.

### Outcome Variable

Dysentery: the presence of bloody diarrhea (as defined by the respondent, the child’s mother). Dysentery was asked by DHS to the mothers whether there was any blood in the stools of those children who had diarrhea at any time during the two weeks preceding the interview (0 = no; 1 = yes).

### Independent Variables

We divided the variables according to the data source: level-1 variables included the child, mother and household characteristics, and level-2 variables included country data. At first, we considered three levels of analysis –child, household, and country-but most households had only one child under the age of five, so it was decided to include only one child per household, the youngest, and conduct a two-level analysis.

#### Level-1 data, children

age of the child in months, sex, possession of the health card, immunization defined as the completeness of WHO schedule, duration of breath feeding in months, birthweight, twin pregnancy, cesarean section, bottle feeding with a nipple. Even though it is not ideal to include a birthweight-missing indicator, taking into consideration that 46% of the children did not have this information, imputation was ruled out.

#### Level −1 data, mother

age in years, education o educational attainment, current employment, marital status, total children and partner age

#### Level −1 data, mother’s pregnancy

planned pregnancy, antenatal visits.

#### Level −1 data, household

household members defined as the number of people living in the same home, place of residence, the age of household head, main floor material, years living at the house, sanitation score based upon water source and waste disposal and wealth index calculated by the DHS considering income, possessions, and quality of life [12].

#### Level-2 data, country

Country wealth coded as a set of dummy variables: Low income (1 = gross domestic product per capita (GDPpc) of US$1,025 or less), Lower middle income (1 = GDPpc between US$1,026 and US$4,035), Upper middle income (1 = GDPpc between $4,036 and $12,475), and High income (1 = GDPpc of $12,476 or more) [15]. Inequality based on the Gini coefficient (1 = top 25 % unequal countries and 0 = more equal countries) and health expenditure coded as a set of dummy variables based on the percentage of GDP expended on health. Low health expenditure (1 = 5 % or less), Middle health expenditure (1 = between 5.1 and 10 %), High health expenditure (1 = more than 10 %). We considered in the initial models country homicide rates and total country population, but these variables were omitted in the final models because of their lack of association with dysentery and their negative effects on the model’s validity, measured using residual files and reliability estimates.

### Statistical Analysis

The analysis was conducted considering known factors associated with dysentery and the country characteristics studied. Multilevel analyses were preferred because the hierarchical nature of the data violated the principles of independence and homogeneity required for a single-level analysis [13].

The statistical analysis was performed using SPSS 20.0 (IBM) and HLM 7 (Scientific Software International, Inc.), as follows: 1) we merged the individual datasets of the 40 countries, 2) we filter out the database following the inclusion and exclusion criteria explained above obtaining information from 33 countries, 3) we calculated descriptive statistics for categorical (proportions) and numerical variables (mean, standard deviation, minimum and maximum values), 4) we obtained bivariate odds ratios using hierarchical linear modeling logistic regressions of dysentery in all of the studied variables, and 5) we generated multivariable models for dysentery using hierarchical linear modeling. Stepwise multilevel logistic regression equations were estimated. Individual, family and household factors were included as possible predictors of dysentery, and differences were deemed to be significant with P-value less than 0.05. The large sample size allowed us to find small differences with narrow 95% confidence intervals. Finally, multilevel modeling was used to explore the association of country characteristics with dysentery (between countries associations) adjusting for individual, family and household predictors of the condition (within countries associations) [14,15]. Full maximum likelihood was used to fit the models. Random effects were estimated only for indicators with variations between groups that could be explained by the studied variables, allowing the coefficients to vary across groups. Those level-1 indicators were centered on the country mean to avoid the problem of co-linearity. All other variables, as well as the neighborhood variables, were centered on the grand mean and we constrained their variance. Level 2 country data variables were dichotomized and analyzed into 50% higher and 50% lower. The final model can be seen in Table 4. We have calculated median odd ratios (MORs) and intra-class correlations (ICC) for the models, as well as 80% interval odds ratios (IORs) for the country level variables [16,17].

We tested bivariate interactions by multiplying duration of breastfeeding and maternal education, duration of breastfeeding and maternal employment, immunization and maternal education, and wealth index and immunization to determine if an interaction was present.

Within countries, weights provided by the DHS for children under 5 years old were employed in the analysis for the level-1 data. They were adjusted to the survey design. Post-stratification was incorporated as a weight adjustment. The adjusted weights were used in all of the analyses. For level-2 data, between countries, weights were created and used in the analysis for each country accounting for their population.

Regression analyses considering the DHS year of survey were performed to assure that the results were not biased by the different time lapses the surveys took place at each country.

Macro International provided the datasets from the 41 countries included in this study. The study was based on secondary sources without identifying information about individual participants. It was given approval by the institutional review board, Comité de Ética en Investigación, Universidad del Rosario.

## Results

The sample comprised 38,762 children under 5 years from 33 developing countries, who presented diarrhea during the last two weeks before the interview, according to the mother’s report. The descriptive features of the evaluated population are shown in Table 1. They are separated from children who did and did not present dysentery in their last episode of diarrhea.

**Table 1.** Descriptive statistics from children under five years old from 33 countries, 2004– 2010 who reported diarrhea. Proportions for categorical variables and median/interquartile range for numerical variables.

| **Table 1.** Descriptive statistics from children under five years old from 33 countries, 2004–2010 who reported diarrhea. Proportions for categorical variables and median/interquartile range for numerical variables. |  |  |
| --- | --- | --- |
| **Variables** | **NO Dysentery** | **Dysentery** |
| **Children** |  |  |
| **Dysentery % (n)** | 85.25 (33,048) | 14.74 (5,714) |
| **Age in months median (IQR)** | 19 (22) | 24 (22) |
| **Female sex % (n)** | 47.32 (15,639) | 47.67 (2,724) |
| **Possession of health card % (n)** | 87.82 (29,026) | 85.35 (4,877) |
| **Complete immunization schedule % (n)** | 63.72 (21,060) | 55.98 (3,199) |
| **Planned pregnancy % (n)** | 57.17 (18,895) | 53.02 (3,030) |
| **Antenatal visits % (n)** |  |  |
| None | 9.47 (3,131) | 11.06 (632) |
| 1-2 | 10.02 (3,312) | 9.53 (545) |
| 3-10 | 59.77 (19,753) | 56.09 (3,205) |
| More than 10 | 4.16 (1,376) | 2.55 (146) |
| Missing information | 16.56 (5,476) | 20.75 (1,186) |
| **Birth weight % (n)** |  |  |
| Low | 6.25 (2,066) | 5.19 (297) |
| Normal | 48.17 (15,921) | 40.47 (2,313) |
| Not weighed or not remember | 45.057(15,061) | 54.32 (3,104) |
| **Cesarean delivery % (n)** | 11.08 (3,665) | 8.76 (501) |
| **Duration of breastfeeding (months) median (IQR)** | 13 (10) | 15 (10) |
| **Still breastfeeding % (n)** | 62.07 (20,514) | 56.10 (3,206) |
| **Drinking from a bottle with a nipple % (n)** | 22.44 (7,418) | 17.72 (1,013) |
| **Twin pregnancy % (n)** | 1.99 (659) | 1.90 (109) |
| **Mother** |  |  |
| **Age (years) median (IQR)** | 27 (10) | 28 (11) |
| **Educational attainment % (n)** |  |  |
| *No education* | 24.23 (8,010) | 30.36 (1,735) |
| *Elementary* | 39.38 (13,017) | 44.50 (2,543) |
| *High school* | 30.24 (9,995) | 21.92 (1,253) |
| *Superior* | 6.12 (2,025) | 3.18 (182) |
| **Mother's Employment % (n)** | 47.76 (15,787) | 57.47 (3,284) |
| **Marital status % (n)** |  |  |
| *Single* | 4.27 (1,669) | 4.48 (256) |
| *Married* | 88.34 (29,195) | 87.73 (5,013) |
| *Divorced separated or widow* | 7.38 (2,440) | 7.78 (445) |
| **Total children ever born median (IQR)** | 3 (3) | 3 (3) |
| **Partner age (years) % (n)** |  |  |
| *Under 20* | 0.58 (193) | 0.36 (21) |
| *Between 20 and 40* | 70.77 (23,391) | 65.68 (3,753) |
| *Above 40* | 16.43 (5,430) | 20.86 (1,192) |
| *Missing information* | 12.20 (4,034) | 13.09 (748) |
| **Household** |  |  |
| **Number of household members median (IQR)** | 6 (4) | 6 (3) |
| **Male-headed households % (n)** | 82.17 (27,157) | 80.71 (4,612) |
| **Age of head of household median (IQR)** | 36 (18) | 37 (17) |
| **Who is the head of household % (n)** |  |  |
| *Mother* | 9.61 (3,179) | 11.37 (650) |
| *Husband* | 64.64 (21,365) | 66.22 (3,784) |
| *Other relative* | 25.73 (8,504) | 22.40 (1,280) |
| **Urban residence % (n)** | 65.20 (21,548) | 73.90 (4,223) |
| **Floor material % (n)** |  |  |
| *Soil or sand* | 44.65 (14,758) | 56.72 (3,241) |
| *Wood* | 11.51 (3,804) | 11.95 (683) |
| *Finished floor* | 43.83 (14,486) | 31.32 (1,790) |
| **Years living at house % (n)** |  |  |
| *Under one* | 4.03 (1,333) | 3.85 (220) |
| *Between one and five* | 22.64 (7,485) | 21.21 (1,212) |
| *More than five* | 62.34 (20,604) | 66.13 (3,779) |
| *Missing information* | 10.97 (3,626) | 8.80 (503) |
| **Inadequate sanitation score % (n)** | 16.70 (5,520) | 23.27 (1,330) |
| **Wealth index % (n)** |  |  |
| *Poorest* | 27.78 (9,182) | 37.45 (2,140) |
| *Poorer* | 22.82 (7,543) | 25.28 (1,445) |
| *Middle* | 20.42 (6,749) | 18.34 (1,048) |
| *Richer* | 16.75 (5,537) | 12.58 (719) |
| *Richest* | 12.21 (4,037) | 6.33 (362) |

The prevalence of dysentery was 14.74%. The median age of the children who had dysentery was 24 months, and the median age among the children who did not present it was 19 months. In addition, only 56% of the children who had dysentery had their immunization schedule completed, in contrast with children who did not present dysentery who accounted for 63%, approximately.

Nearly half of the children who did not have dysentery had normal weight (48%). However, this percentage was lower among the group of children who did present the illness (41%). Moreover, this last group of children was breastfed a median of 15 months, in comparison with the group of children who did not present dysentery and was breastfed a median of 13 months.

Approximately, 24% of the mothers did not have any level of education in the group of children who had present dysentery. Nevertheless, this value was higher in the group of the mothers of children who did present dysentery (30%). Additionally, more than half of this last group of mothers was employed (57%). Finally, the higher the wealth index was, the lower its percentage became in both groups of children.

Furthermore, the prevalence of dysentery in each of the evaluated countries is shown in Figure 1. The Republic of Liberia presented the highest prevalence of dysentery in the whole group of countries, and Colombia had the highest one among the evaluated Latin-American countries.

**Figure 1.**
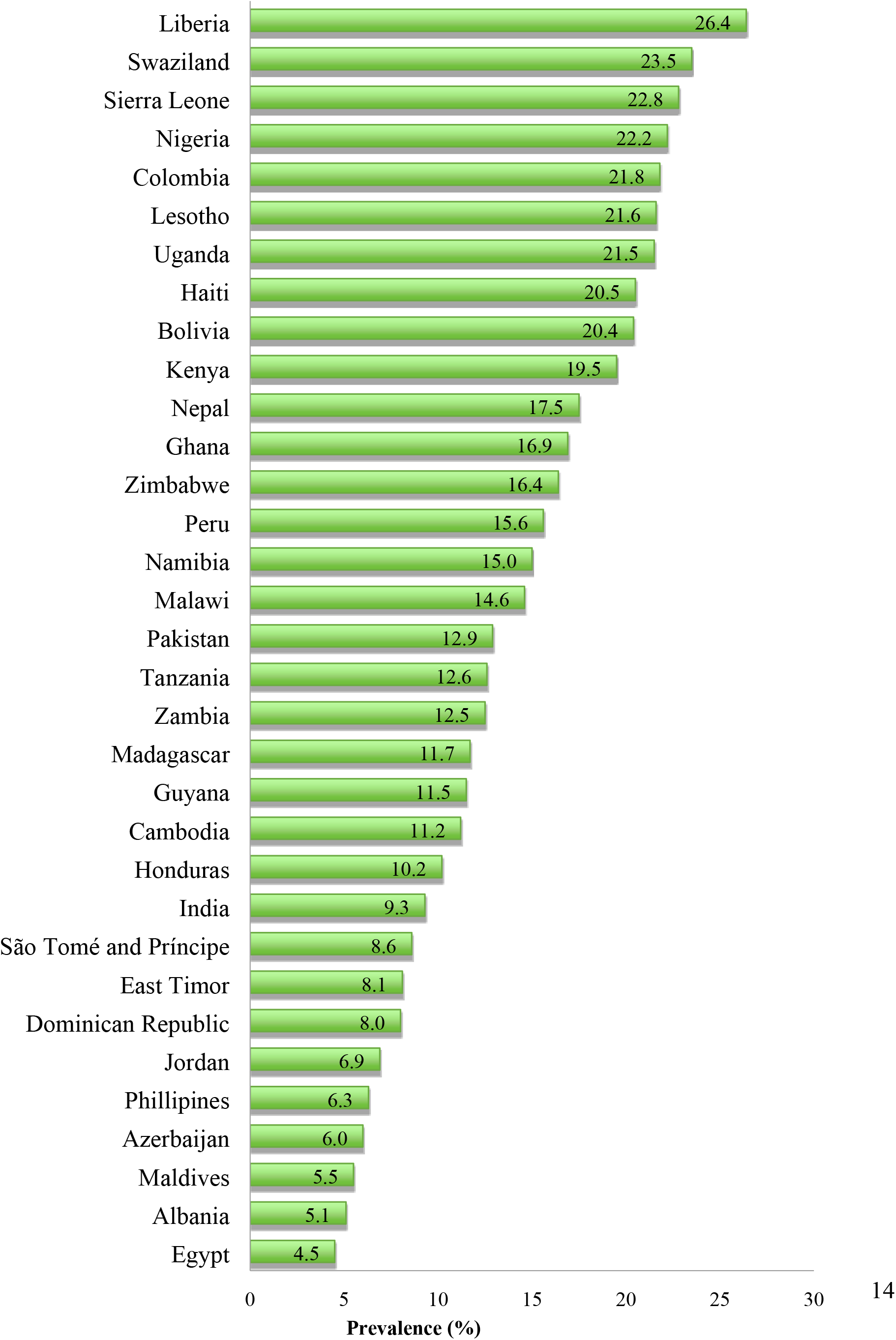
Prevalence of dysentery in evaluated countries

The characteristics of diarrhea in both groups of children are shown in Table 2. Approximately, 58% of children who had dysentery also had fever. Likewise, this group of children received almost 55% and 27% of oral rehydration and antibiotic therapy, respectively. However, they did receive lower amounts of liquids and solids during illness (61% and 38%, respectively), than the group of children who did not present dysentery (67% and 43%, respectively).

**Table 2.**
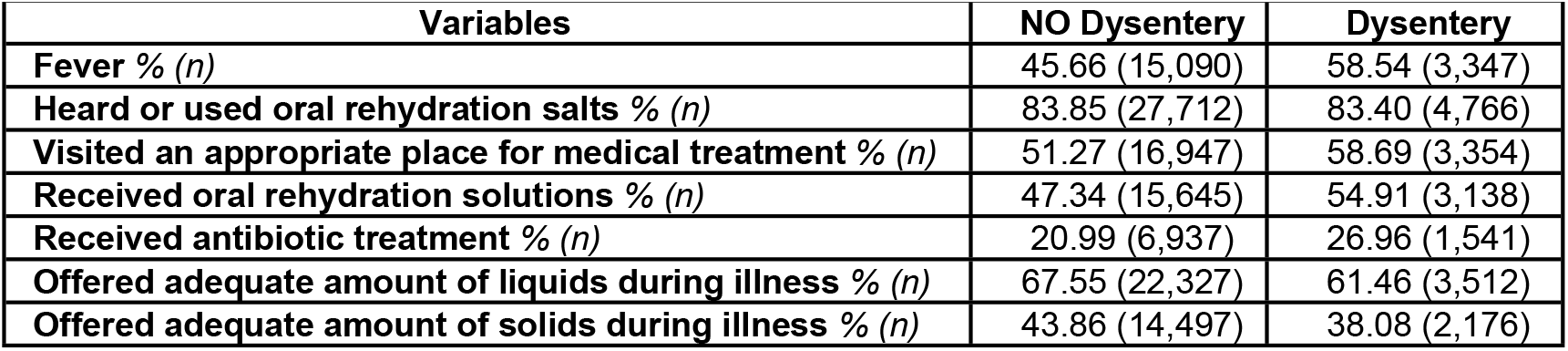
Signs, symptoms and characteristics related to dysentery from children under five years from 33 countries, 2004–2010.

|  |  |  |
| --- | --- | --- |
| **Variables** | **NO Dysentery** | **Dysentery** |
| **Fever % (n)** | 45.66 (15,090) | 58.54 (3,347) |
| **Heard or used oral rehydration salts % (n)** | 83.85 (27,712) | 83.40 (4,766) |
| **Visited an appropriate place for medical treatment % (n)** | 51.27 (16,947) | 58.69 (3,354) |
| **Received oral rehydration solutions % (n)** | 47.34 (15,645) | 54.91 (3,138) |
| **Received antibiotic treatment % (n)** | 20.99 (6,937) | 26.96 (1,541) |
| **Offered adequate amount of liquids during illness % (n)** | 67.55 (22,327) | 61.46 (3,512) |
| **Offered adequate amount of solids during illness % (n)** | 43.86 (14,497) | 38.08 (2,176) |

Table 3 shows the results of the bivariate regressions among dysentery and each one of the others included variables.

**Table 3.** Bivariate regressions of dysentery on independent variables

| **Table 3.** Bivariate regressions of dysentery on independent variables |  |  |
| --- | --- | --- |
| **Variables** | **OR** | **P- value** |
| **Children** |  |  |
| Age (months) | 1.01 | 0.000 |
| Female sex | 1.01 | 0.624 |
| Possession of health card | 0.81 | 0.000 |
| Complete immunization schedule | 0.72 | 0.000 |
| Planned pregnancy | 0.85 | 0.000 |
| None antenatal visit | 1.19 | 0.000 |
| 1-2 antenatal visits | 0.95 | 0.259 |
| 3-10 antenatal visits | 0.86 | 0.000 |
| More than 10 antenatal visits | 0.60 | 0.000 |
| Missing information about antenatal visits | 1.32 | 0.000 |
| Low weight at birth | 0.82 | 0.002 |
| Normal weight at birth | 0.73 | 0.000 |
| Not weighted at birth or not remember | 1.42 | 0.000 |
| Cesarean delivery | 0.77 | 0.000 |
| Duration of breastfeeding (months) | 1.03 | 0.000 |
| Still breastfeeding | 0.78 | 0.000 |
| Drinking from a bottle with a nipple | 0.75 | 0.000 |
| Twin pregnancy | 0.96 | 0.665 |
| **Mother** |  |  |
| Age (years) | 1.02 | 0.000 |
| No education | 1.36 | 0.000 |
| Elementary | 1.23 | 0.000 |
| High school | 0.65 | 0.000 |
| Superior | 0.50 | 0.000 |
| Employed | 1.48 | 0.000 |
| Single | 1.50 | 0.482 |
| Married | 0.94 | 0.187 |
| Divorced | 1.06 | 0.282 |
| Total children ever born | 1.10 | 0.000 |
| Partner under 20 years old | 0.63 | 0.041 |
| Partner between 20 and 40 years old | 0.79 | 0.000 |
| Partner above 40 years old | 1.34 | 0.000 |
| Missing information about partner age | 1.08 | 0.061 |
| **Household** |  |  |
| Number of household members | 1.03 | 0.000 |
| Female household head | 1.10 | 0.008 |
| Age of head of household | 1.00 | 0.760 |
| Mother household head | 1.21 | 0.000 |
| Husband household head | 1.07 | 0.021 |
| Other relative household head | 0.83 | 0.000 |
| Rural residence | 0.66 | 0.000 |
| Soil or sand | 1.62 | 0.000 |
| Wood | 1.04 | 0.334 |
| Finish floor | 0.59 | 0.000 |
| Poorest | 1.56 | 0.000 |
| Poorer | 1.15 | 0.000 |
| Middle wealth index | 0.88 | 0.000 |
| Richer | 0.72 | 0.000 |
| Richest | 0.49 | 0.000 |
| Adequate sanitation score | 0.66 | 0.000 |
| **Illness** |  |  |
| Fever | 1.68 | 0.000 |
| Heard or used oral rehydration salts | 0.97 | 0.400 |
| Visited an appropriate place for medical treatment | 1.35 | 0.000 |
| Received oral rehydration solutions during illness | 1.36 | 0.000 |
| Received antibiotic treatment during illness | 1.39 | 0.000 |
| Offered adequate amount of liquids during illness | 0.77 | 0.000 |
| Offered adequate amount of solids during illness | 0.79 | 0.000 |

Results of logistic regression among dysentery and the country level characteristics are shown in Table 4. After controlling for each of the child, mother, house and country level variables, we found that related to dysentery, GDP per-capita and health expenditure were negatively associated, and GINI index was positively associated.

**Table 4.** Multivariate multilevel logistic regressions for dysentery and country level characteristics.

|  |  |  |  |
| --- | --- | --- | --- |
| **Variables** | **Final model** |  |  |
|  | **OR** | **CI** | **p-value** |
| *Child sex* | 1.03 | (0.96-1.11) | 0.337 |
| *Child age* | 0.99 | (0.99-1.00) | <0.001 |
| *Vaccination* | 0.88 | (0.81-0.96) | 0.003 |
| *Normal weight at born* | 0.95 | (0.87-1.04) | 0.276 |
| *Duration of breastfeeding* | 0.81 | (0.75-0.89) | <0.001 |
| *Mother's age* | 1.01 | (1.00-1.01) | 0.002 |
| Mother's education |  |  |  |
| *No education* | 1.25 | (1.84-1.60) | 0.049 |
| *Elementary level education* | 1.24 | (0.98-1.57) | 0.065 |
| *High school level education* | 1.03 | (0.82-1.29) | 0.798 |
| Superior level education | Comparison category |  |  |
| *Employed mother* | 1.11 | (1.02-1.20) | 0.009 |
| *Married* | 1.07 | (0.95-1.20) | 0.242 |
| *Number household members* | 1.02 | (1.01-1.03) | <0.001 |
| *Type of residence urban* | 0.87 | (0.79-0.97) | 0.015 |
| Household wealth |  |  |  |
| *Poorest* | 2.07 | (1.72-2.49) | <0.001 |
| *Poorer* | 1.78 | (1.49-2.13) | <0.001 |
| *Middle* | 1.49 | (1.25-1.78) | <0.001 |
| *Richer* | 1.38 | (1.16-1.65) | <0.001 |
| *Richest* | Comparison category |  |  |
| INTERCEPT | 0.14 | (0.12-0.16) | <0.001 |
| GDP per capita | 0.75 | (0.71-0.78) | 0.003 |
| Gini index | 1.23 | (1.19-1.28) | <0.001 |
| Health expenditure | 0.85 | (0.80-0.89) | <0.001 |
| **<u>Random effects</u>** | ***Standard deviation*** | ***Variance component*** | ***p-value*** |
| Intercept | 0.34 | 0.12 | <0.001 |
| **<u>Reliability</u>** |  |  |  |
| Intercept | 87.8% |  |  |

At the same time, this last model showed that child age, mother age, employed mother, and the number of household members have significant positive associations with having dysentery (p-value less than 0.05). On the other hand, complete immunization schedule, duration of breastfeeding, and the type of residence have significant negative associations with having the illness (p-value less than 0.05).

Simultaneously, each one of the categories of wealth index showed a significant association with dysentery (p-value less than 0.001). And it is possible to appreciate that the richer a person is, the lower its odds ratios becomes

## Discussion

In this study, 14.7%, of the 38,762 children under-five years who suffered from diarrhea, had dysentery. This is consistent with the following data found. Diarrheal diseases remain among the most common causes of mortality and morbidity in children, particularly in low and middle-income countries. In 2013, of the 6.3 million children worldwide who died before their fifth birthday about 7.94% died from diarrhea (18). In a study in North Ethiopia with 241 participants, the overall prevalence was 13.3% (19).

Similar to previous studies, we observed a positive correlation between dysentery and mother’s age, no education of the mother, employed mother, the number of household members and poverty [21–23], as well as a negative correlation between dysentery and child age, vaccination, duration of breastfeeding and urban residence [24,25].

It is worth mentioning that the household wealth showed a gradient of association with dysentery that changed depending on the wealth category. This confirms what was found by Chompook et al. in Thailand, [26] the poorer you are, the more likely to get dysentery.

The country characteristics studied showed association with dysentery. A negative correlation of GDP per capita and child dysentery was observed, which means that the lack of economic production is associated with the health of children at the country level. This finding is consistent with other epidemiological studies, which present how the lack of economic resources at the country level is associated with the decrease and absence of opportunities and in turn with impaired health of the population [27,28].

The Gini index showed a positive correlation with dysentery. The degree of inequality in the distribution of income is related to the health of children at the country level. Inequalities create the sense of unfairness and feelings of injustice and discrimination in the disadvantaged group, due to the difference in the opportunities offered [29]. These feelings have the potential to undermine the wellbeing of children and their families.

Health expenditure also showed a positive correlation to dysentery. It is likely that when a country invests their money in health, it is giving its children the potential to be healthier [30].

## Limitations

Even though DHS has significant and well-known advantages related to quality, comparability, and representativeness of the information, our study presents some significant limitations.

First, due to its cross-sectional nature, it is not possible to establish any causal relationship among studied variables. Second, as data was collected exclusively from mothers, who are supposed to be the best relators about their child’ s history, related bias are likely to be found. Third, while DHS questionnaires are not executed simultaneously in every country and social conditions tend to change over time, some differences could be expected. However, our results did not change after controlling for year of survey. Fourth, the variable definitions are limited by the established methodology of the DHS team. Finally, the resulting large sample size contributes to an over-power analysis that could detect minimal effect sizes, and these could mean slight biases in the sampling process.

## Conclusion

This study explored, by using a multilevel analysis, the association between per capita GDP, income inequalities (Gini coefficient), health expenditure and dysentery, in a multinational population adjusted by individual, maternal and household characteristics. We found that some factors like older age of the child and their mother, an unemployed mother, lower number of household members, higher wealth index, and higher Gini index are protective factors against dysentery; while, lower GDP per-capita, incomplete immunization schedule, lower duration of breastfeeding and rural residence are risk factors against the same illness.

Additionally, as in a previous study, health expenditure does not appear to take part in developing dysentery. However, per capita GDP and Gini coefficient keep showing an important involvement with progressing from acute diarrhea to dysentery. Due to this, and in order to diminish the consequences of this morbid presentation of acute diarrhea, countries weigh up ways of improving their per capita GDP and diminishing inequalities. Studying associated factors with developing dysentery during an episode of acute diarrhea could be the base upon which we can diminish mortality from this illness.

## Supporting Information

**S1 Fig. Prevalence of dysentery in evaluated countries.**

**S1 Table. Descriptive statistics from children under five years from 33 countries, 2004–2010 who reported diarrhea.** Proportions for categorical variables and median/interquartile range for numerical variables.

**S2 Table. Signs, symptoms and characteristics related to dysentery from children under five years from 33 countries, 2004–2010.**

**S3 Table. Bivariate regressions of dysentery on independent variables**

**S4 Table. Multivariate multilevel logistic regressions for dysentery and country level characteristics.**

| Country | Year | Country | Year |
| --- | --- | --- | --- |
| Albania | 2008–2009 | Malawi | 2010 |
| Azerbaijan | 2006 | Maldives | 2009 |
| Bangladesh | 2007 | Mali | 2006 |
| Benin | 2006 | Namibia | 2006–2007 |
| Bolivia | 2008 | Nepal | 2006 |
| Cambodia | 2010 | Niger | 2006 |
| Colombia | 2010 | Nigeria | 2008 |
| Congo | 2005 | Pakistan | 2006–2007 |
| Egypt | 2005–2006 | Peru | 2004–2008 |
| Philippines | 2008 | Democratic Republic of Congo | 2007 |
| Ghana | 2008 | Dominican Republic | 2007 |
| Guyana | 2009 | São Tomé é Príncipe | 2008–2009 |
| Haiti | 2005–2006 | Sierra Leone | 2008 |
| Honduras | 2005–2006 | Swaziland | 2006– 2007 |
| India | 2005-2006 | Tanzania | 2010 |
| Indonesia | 2007 | East Timor | 2009–2010 |
| Jordan | 2007 | Ukraine | 2007 |
| Kenya | 2008–2009 | Uganda | 2006 |
| Lesotho | 2009 | Zambia | 2007 |
| Liberia | 2007 | Zimbabwe | 2005–2006 |
| Madagascar | 2008–2009 |  |  |

